# Diverse demographic histories in a guild of hymenopteran parasitoids

**DOI:** 10.1101/2019.12.20.884403

**Authors:** William Walton, Graham N Stone, Konrad Lohse

## Abstract

Signatures of changes in population size have been detected in genome-wide variation in many species. However, the causes of such changes and the extent to which they are shared across co-distributed species remain poorly understood. During Pleistocene glacial maxima, many temperate European species were confined to southern refugia. While vicariance and range expansion processes associated with glacial cycles have been widely studied, little is known about the demographic history of refugial populations, and the extent and causes of demographic variation among codistributed species. We used whole genome sequence data to reconstruct and compare demographic histories during the Quaternary for Iberian refuge populations in a single ecological guild (seven species of chalcid parasitoid wasps associated with oak cynipid galls). We find support for large changes in effective population size (*N*_*e*_) through the Pleistocene that coincide with major climate change events. However, there is little evidence that the timing, direction and magnitude of demographic change are shared across species, suggesting that demographic histories are largely idiosyncratic. Our results are compatible with the idea that specialist parasitoids attacking a narrow range of hosts experience greater fluctuations in *N*_*e*_ than generalists.

## Introduction

Natural populations change in size and distribution in response to biotic and abiotic factors over both ecological and evolutionary timescales. Given that population size is a fundamental parameter in the evolutionary process, there is considerable interest in using genetic variation in modern samples to reconstruct the demographic history of populations (see Beichman et al., 2018, for a recent review). A key question is to what extent demographic history has been shaped by past climatic events and, if so, whether such responses are concordant across taxa, or vary with species traits.

During Pleistocene glacial maxima, the ranges of many temperate European plant and animal taxa were restricted to southern refugia, primarily in Iberia, Italy and the Balkans (Hewitt, 2000; Feliner, 2011; Hofreiter and Stewart, 2009). Modern refugial populations of such taxa commonly harbour higher genetic diversity than northern populations founded by postglacial range expansion (e.g. Hewitt, 2000; Rokas et al., 2003; Tison et al., 2014; Vitales et al., 2016) (but see Comps et al., 2001). However, little is known about how refugial population sizes varied through glacial cycles, and the extent to which such variation is concordant across co-distributed species. One hypothesis is that contraction of suitable habitat during glacial maxima reduced population sizes simultaneously across suites of ecologically interacting species. Alternatively, the long term size of refugial populations may be shaped by gene flow both within and between major refugia during brief warmer periods within glacial maxima (Cooper et al., 2015; Hofreiter and Stewart, 2009; Tallavaara et al., 2015). Persistent spatial structure within refugia has been discovered in some species, leading to a hypothesis of ‘refugia within refugia’ (Gómez and Lunt, 2007; Feliner, 2011). In this case, the long term genetic diversity of a species is predicted to be a function of its dispersal ability. We might also expect ecologically specialist species (whose biology is critically dependent on interactions with a small number of other taxa) to experience greater or more frequent demographic changes than generalists (whose demography is less tightly coupled to abundance of any specific interaction) (Rand and Tscharntke, 2007; Östergård and Ehrlén, 2005). While specialists have been shown to be more vulnerable than generalists to habitat fragmentation and climate change in taxa ranging from parasites and parasitoids (Cizauskas et al., 2017; Rand and Tscharntke, 2007) to fungi (Nordén et al., 2013) and mammalian predators (Janecka et al., 2016), few studies have investigated whether specialists have experienced stronger or more frequent *N*_*e*_ changes than generalists over evolutionary timescales (Mackintosh et al., 2019). A third alternative is associated with a pattern of longitudinal range expansion across Europe seen in some taxa (including the system studied here) through the Pleistocene (Stone et al., 2012; Bunnefeld et al., 2018). For refugial populations that formed during the Pleistocene, we might expect signatures of past increases in population size. Here we discriminate between these alternative hypotheses for Iberian refuge populations of multiple species in a single ecological guild - chalcid parasitoid wasps associated with herbivorous cynipid gall wasp hosts on oak.

Oak gall wasp communities are multitrophic systems comprising oak (*Quercus*) host plants, cynipid gall wasp herbivores and chalcid parasitoid natural enemies (Stone et al., 2002). The latter are obligate specialists of cynipid galls, allowing the guild of parasitoids to be considered in ecological isolation. Many of the component species show broad longitudinal distributions, extending from Iberia in the West to Iran in the East, and all species studied to date are genetically structured into major southern refuge populations (Stone et al., 2012, 2017; Nicholls et al., 2010, 2012; Lohse et al., 2010, 2012; Petit et al., 2002). Bunnefeld et al. (2018) showed that refugia in the Balkans and Iberia were colonized primarily through westwards range expansion from an eastern origin one or more glacial cycles in the past. Iberian populations have lower genetic diversity than more eastern refugia, show little evidence of population structure (Rokas et al., 2003; Stone et al., 2007; Nicholls et al., 2010) (for an exception see Rokas et al. (2001), and are the source of most post-colonisation gene flow (Stone et al., 2017; Bunnefeld et al., 2018). Thus for most species in this community the Iberian refuge population represents the end-point of a longitudinal expansion history.

Here we use whole genome sequence (WGS) data to quantify and compare the demographic histories of Iberian refuge populations in seven chalcid parasitoid species in the oak gall wasp community. Our sampling targets males (five from Iberia, one from the Balkans in each species), whose haploid genome facilitates analysis (Bunnefeld et al., 2018; Hearn et al., 2014). WGS data allow for powerful demographic inference even for non-model organisms. However, given fragmented reference genomes and limited sample sizes available for these species, inference methods must be chosen carefully and, ideally, should use linkage as well as allele frequency information. In particular, inference based only on the site frequency spectrum (SFS) (e.g. Gutenkunst et al., 2009) requires larger samples for accurate inference of recent population history. We investigate demographic history using two approaches that use the signal contained in genome-wide variation of a small sample of individuals in different ways (Bunnefeld et al., 2018; Lohse et al., 2011; Li and Durbin, 2011): we use a parametric maximum-composite likelihood (MCL) method based on a blockwise summary of sequence variation (Lohse et al., 2011) (hereafter termed the ‘blockwise method’) to fit a model of a single instantaneous step change in *N*_*e*_. Using analytic likelihood calculations to fit such fully specified but minimally complex histories utilises both frequency and linkage information and facilitates comparison between species for the timing and magnitude of *N*_*e*_ change. We also applied a non-parametric method, the Pairwise Sequentially Markovian Coalescent (PSMC) (Li and Durbin, 2011), based on the distribution of pairwise differences in minimal samples of two haploid genomes. PSMC generates a more resolved picture of *N*_*e*_ change through time, including an estimate of the time at which populations have diverged. As the two methods use different data properties and sampling strategies, we can have high confidence in inferences supported by both. Both methods have contrasting limitations: while the blockwise method – by design – cannot detect gradual *N*_*e*_ changes or resolve population histories involving multiple changes, PSMC is known to smooth out very sudden changes. PSMC is based on fewer samples which contain less information (i.e. fewer coalescence events) about the recent *N*_*e*_ (Li and Durbin, 2011). The two methods therefore complement each other and together provide a comprehensive picture of population history.

We address the following questions: *(i)* What signatures of *N*_*e*_ change, if any, are present for each species? *(ii)* To what extent are the direction and timing of changes in *N*_*e*_ concordant across species? Do species show evidence for simultaneous *N*_*e*_ change suggesting concordant responses to a shared underlying driver such as climate change, or are their demographic histories largely idiosyncratic? *(iii)* Do major demographic changes coincide with specific glacial or interglacial periods in the Pleistocene? *(iv)* Did inferred demographic changes occur after the colonisation of Iberia, or are they shared with other refugial populations, suggesting that they took place in a shared ancestral population prior to the colonisation of Iberia?

## Materials and Methods

### Samples and sequencing

We analysed whole genome Illumina paired end resequencing data for six (haploid) male individuals in each of seven species of chalcid parasitoids (spanning five families): *Megastigmus dorsalis* and *M. stigmatizans* (Megastigmidae), *Torymus auratus* (Torymidae), *Ormyrus nitidulus* and *O. pomaceus* (Ormyridae), *Eurytoma brunniventris* (Eurytomidae) and *Cecidostiba fungosa* (Pteromalidae) (Table S1). The *M. dorsalis* individuals correspond to cryptic species 1 for this complex, as defined by Nicholls et al. (2010). For each species, we sampled five individuals from Iberia (the focal refugial population) and one from Hungary (the Balkan refugium). Data for the Hungarian and two Iberian samples of each species were generated previously by Bunnefeld et al. (2018). We generated analogous data for the remaining three Iberian individuals using the protocols described by Bunnefeld et al. (2018). DNA was extracted from individual wasps using the Qiagen DNeasy kit. Individual Nextera genomic libraries were generated and sequenced on an Illumina HiSeq 2000 by the NERC Edinburgh Genomics facility, UK. Raw reads were deposited at the SRA (PRJEB20883). Mean coverage per haploid individual ranged from 3.5x to 18.5x. Raw reads were mapped back to reference genomes assembled by Bunnefeld et al. (2018) using *BWA* (0.7.15-r1140) (Li and Durbin, 2009), duplicates were marked with *picard* (V2.9.0) MarkDuplicates (broadinstitute.github.io/picard/), variants were called using *Freebayes* (v1.1.0-3-g961e5f3) (https://github.com/ekg/freebayes) with a minimum base quality of 10 and a minimum mapping quality of 20 (see Bunnefeld et al. (2018) for further details).

### Inferring step changes in population size

We fitted a model of a single instantaneous step change in *N*_*e*_ using the framework for blockwise likelihood calculations developed by Lohse et al. (2011). The model includes three parameters: the scaled mutation rate *θ* = 4*N*0*µ× l* (where *N*0 is the current *N*_*e*_ and *l* is block length); *T*, the time of *N*_*e*_ change measured in 2*N*_*e*_ generations; and *λ* = *N*_0_*/N*_1_, the relative magnitude of the *N*_*e*_ change, (where *N*_1_ denotes the *N*_*e*_ prior to the step change).

Following Bunnefeld et al. (2015), we summarized sequence variation in short blocks of a fixed length *l* by the (folded) blockwise site frequency spectrum (bSFS). Given our sampling scheme of *n* = 5 haploid males, the bSFS consists of counts of two types of variants: those for which the minor allele occurs once or twice in the sample. The probability of observing a particular set of mutations in a block can be computed analytically as a higher-order derivative of the generating function of genealogies (Lohse et al., 2011). The product of probabilities of bSFS configurations across blocks can be interpreted as the composite likelihood (*CL*) of the model. We maximised *lnCL* in *Mathematica* (Wolfram Research, 2016) using the function *NMaximise* (Supplementary File 1).

To generate blockwise data for each species, we applied the same quality filters used for calling SNPs to all sites, i.e. we identified regions of the genome with a base quality *>*10 and mapping quality *>*20 in each individual from bam files by the CallableLoci walker of *GATK* (v3.4) (Van der Auwera et al., 2013). Only regions meeting these criteria in all five Iberian individuals were included in further analyses. Custom scripts were used to partition the data into blocks of a fixed length *l* of callable sites. *l* was chosen to be inversely proportional to the pairwise genetic diversity of each species (Table 2), such that blocks contained on average two pairwise differences. This ensures that the information content per block as well as any effect of intra-block recombination is consistent across species. Blocks with a physical span (including non-callable sites) of *>* 2*l*, contigs with length *<* 2*l*, and blocks with more than five “None” (uncalled) sites were removed.

We assessed support for a step change in *N*_*e*_ relative to the (nested) null model of constant *N*_*e*_ by generating parametric bootstraps with the coalescent simulator *msprime* (Kelleher et al., 2016). For each species, 100 replicate data sets were simulated under the null model assuming estimates of recombination inferred by Bunnefeld et al. (2018) (Table S3). Each dataset had the same total length as the real data (after filtering) and was partitioned into 5,000 windows of sequences (for computational efficiency). Both a null model of constant *N*_*e*_ and a step change model were fitted to each replicate, and the 95% quantile of the difference in support between models *lnCL*s was compared to that of the real data. Confidence intervals (CIs) of parameter estimates were obtained via an analogous parametric bootstrapping procedure: we simulated 100 datasets with recombination under the inferred step change model and fitted that model to each simulation replicate. CIs were obtained as 2.5% and 97.5% quantiles of the distribution of parameter estimates, and were centred around the point estimates of parameters obtained from the real data.

### Reconstructing ancestral population size with PSMC

The Pairwise Sequentially Markovian Coalescent (PSMC) (Li and Durbin, 2011) was used to infer a history of population size change in each species. PSMC is a non-parametric method that reconstructs a trajectory of past *N*_*e*_ from the density of pairwise differences along the genome via a hidden Markov model in which the hidden states are pairwise coalescence times, the distribution of which is used to estimate *N*_*e*_ in discrete time intervals. In each species, the two Iberian individuals with the greatest average read depth were used as the focal pair. Fastq files were generated using *samtools mpileup* (Li et al., 2009), and regions covered in both individuals were combined into fastq files using *seqtk mergefa* (https://github.com/lh3/seqtk) and converted into PSMC input files. PSMC, by default, discretizes pairwise alignments into blocks of 100bp which are encoded as variant if they contain at least one variant. While this makes analyses of large genomes with low diversity (e.g. humans) computationally efficient, this discretisation is too coarse when considering more diverse genomes where the chance of several pairwise differences occurring in the same 100 bp block is non-negligible (which biases *N*_*e*_ estimates downwards). We investigated the effect of varying block length (100, 50, 25 and 1bp) on *N*_*e*_ inference for the most diverse (*E. brunniventris*, *π* = 0.0071) and the least diverse (*M. stigmatizans π* = 0.00067) species in our set. As expected, population trajectories showed higher *N*_*e*_ and were pushed back in time with smaller block lengths (Figure S1). We chose a block length of 25bp for all analyses which minimizes these biases without excessive computational cost.

We inferred 30 free interval parameters across 64 time intervals (with the option -p “28*2+3+5”). The maximum coalescence time (-t) was set to 5, the initial value of *θ/ρ* (-r) to 1. 100 bootstraps were performed for each run. Times of peak *N*_*e*_ and values of *N*_*e*_ in each time interval were considered to be different between species if there was no overlap in bootstrap replicates. To be able to compare the magnitude of inferred *N*_*e*_ change between PSMC and blockwise analyses, we normalised the maximum *N*_*e*_ inferred by each method by the long-term average *N*_*e*_ estimated simply from *π* (Table 2) as 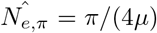 and computed the following measure of *N*_*e*_ change: 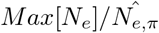. Unlike the size of the step change (*λ*), this measures the maximum *N*_*e*_ relative to the average over the entire history of a sample and is therefore expected to be smaller than *λ*.

### Calibrating the timing of events

Time estimates were converted into years assuming a mutation rate of 3.46 *×* 10^*−9*^ mutations per base per generation estimated for *Drosophila melanogaster* (Keightley et al., 2009). We assumed two generations per year for all species with the exception of *M. stigmatizans* which has a single generation per year (Stone et al., 2012). While this calibration allows comparisons with the estimates obtained by Bunnefeld et al. (2018) (calibrated in the same way), these absolute times are likely underestimates and should be treated with caution (given that we use a spontaneous mutation rate estimate and not all *de novo* mutations are selectively neutral). We stress, however, that comparing the relative timing of demographic events across this set of parasitoids only relies on the assumption of the same per generation mutation rate across species rather than any absolute calibration.

### Cross-population PSMC analyses

To test whether population size changes in each species occurred before or after the divergence between the Iberian and Balkan refuge populations, we compared the PSMC trajectories of Iberian pairs and cross-population pairs (one Iberian and one Hungarian individual). Any divergence between Iberian and the Balkan refuge populations should be visible as a separation between the within-population (Iberian pair) and the cross-population (Iberia-Balkan pair) PSMC trajectories. Specifically, in the absence of post-divergence gene flow we expect *N*_*e*_ estimates for the cross-population pairs to increase from the time of the population split due to the accumulation of genetic differences between the populations. Thus population size changes that occurred in an ancestral population (potentially outside Iberia) should predate the divergence of within and cross-population trajectories. In contrast, we would expect demographic events unique to the Iberian population to happen after divergence of the within- and cross-population PSMC trajectories. We also compared our PSMC divergence time estimates to those made by Bunnefeld et al. (2018) under explicit models of divergence and admixture.

### Potential population structure in Iberia

Population structure within any assumed panmictic population (here, Iberia) can confound inferences of past demography (Gattepaille et al., 2013; Bunnefeld et al., 2015). In particular, *N*_*e*_ estimates may be inflated due to divergence between demes (Gattepaille et al., 2013). Likewise, when samples are taken from the same deme, the (structured) coalescent generates a mixture of very recent (within-deme) and older ancestry resulting from migration between demes (Wakeley, 2008), which can mimic signatures of a bottleneck. To test for population structure within Iberia, we repeated PSMC analyses on every pairwise combination of our five individuals. In a structured population, and given that the sampling locations were widely spaced across Iberia (Table S1), *N*_*e*_ trajectories are expected to differ between pairs of haploid males sampled from the same *versus* different sub-populations (analogous to the within- and between-population analyses involving Iberian and Hungarian samples).

### Sensitivity analyses

The assumption of no recombination within blocks (which is required to make the composite likelihood estimation tractable) potentially leads to biases in parameter estimates. Specifically, recombination may lead to a downward bias of *N*_*e*_ estimates (Wall, 2003; Bunnefeld et al., 2015). We used simulations in *msprime* (Kelleher et al., 2016) to quantify the effect of recombination on parameter estimates (Table S2). One million unlinked blocks of 586bp (corresponding to the block length used for *O. nitidulus*) were simulated under a step change model with different recombination rates and step sizes.

It is well known that both selective sweeps (Smith and Haigh, 1974) and background selection (Charlesworth et al., 1993) affect variation at neutral sites in the genome which, in turn, can bias demographic inference (Ewing and Jensen, 2016; Schrider et al., 2016). To explore the effect of selection on the blockwise analyses, we fitted step change models separately to blockwise data generated from genic and intergenic regions of the *O. nitidulus* genome. Genes were predicted *ab initio* with *Augustus* (Stanke and Morgenstern, 2005). *Nasonia vitripennis*, a chalcid parasitoid (family Pteromalidae), was used as a training set. If selection has a strong effect on demographic inference, estimates of *θ* are expected to be much lower for genic compared to intergenic regions, as most selection tends to reduce diversity both at selective targets and linked regions of the genome (Smith and Haigh, 1974). Similarly, we would expect estimates for the time of the step change, *T*, to be biased towards the present in genic compared to intergenic regions.

## Results

After filtering, the blockwise datasets ranged in total length from 54 to 151 Mb (Table S3). The pairwise alignments used as input for PSMC spanned a total length of 161 to 379 Mb (Table S3). Despite this difference in overall length, which is mainly a due to missing data and the difference in sample size (two *versus* five individuals), estimates of average pairwise diversity, as measured by *π* (Nei, 1972), agreed broadly between the two datasets and were slightly lower for the blockwise data (Table 1) compared to the pairwise alignments (Table 2). Across species, *π* estimates spanned an order of magnitude, from 0.00067 in *M. stigmatizans* to 0.0071 in *E. brunniventris*.

**Table 1:** Maximum composite likelihood estimates (MCLE) of demographic parameters for Iberian populations of seven species of chalcid parasitoid wasps under a model of a single step change. Estimates for the time of *N*_*e*_ change (*T*) are given in thousands of years ago (kya). 95% CIs are shown in brackets. The maximum *N*_*e*_ is scaled relative to *π* as *Max*[*N*_*e*_]/*N*_*e,π*_.

| Species | $\pi$ | Current $N_e \times 10^3$ | Ancestral $N_e \times 10^3$ | $T$ (kya) | Relative $Max[N_e]$ |
| --- | --- | --- | --- | --- | --- |
| *T. auratus* | 0.00509 | 1,142 (1,126-1,157) | 422 (418-425) | 100 (94-106) | 2.14 |
| *M. stigmatizans* | 0.00054 | 18 (14-21) | 61 (60-61) | 10 (9.5-10.4) | 1.55 |
| *M. dorsalis* | 0.00142 | 67 (59-75) | 144 (143-145) | 12.5 (12.0-13.0) | 1.40 |
| *O. nitidulus* | 0.00126 | 47 (45-49) | 194 (192-197) | 33 (31-35) | 2.13 |
| *O. pomaceus* | 0.00230 | 207 (207-208) | 207 n/a | n/a | n/a |
| *E. brunniventris* | 0.00569 | 523 (521-525) | n/a | n/a | n/a |
| *C. fungosa* | 0.00258 | 234 (233-235) | n/a | n/a | n/a |

**Table 2:**
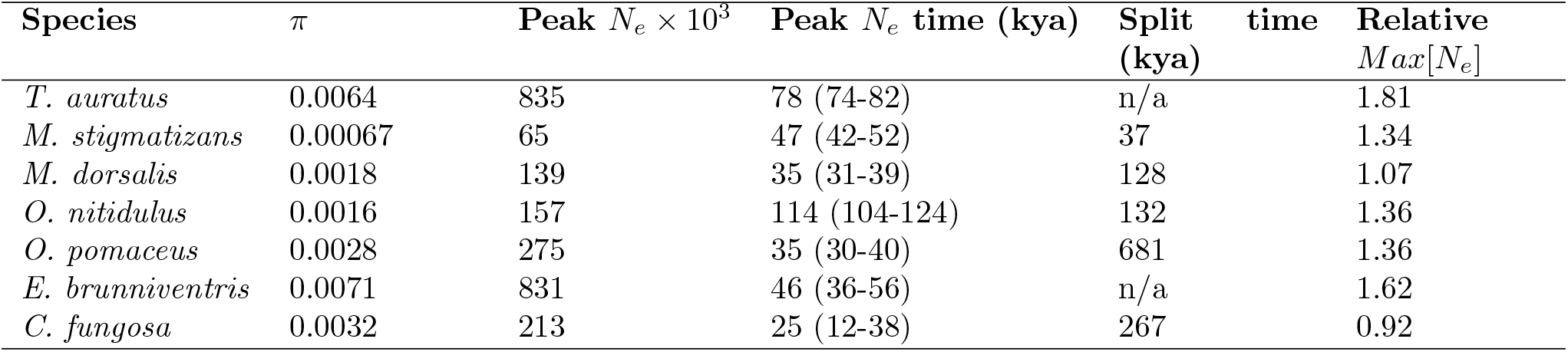
Summaries of PSMC trajectories for Iberian populations of seven species of chalcid parasitoid wasps. Times are given in thousands of years ago (kya). 95 % CIs of peak *N*_*e*_ times are shown in brackets. The maximum *N*_*e*_ is scaled relative to *pi* as *Max*[*N*_*e*_]/*N*_*e,π*_.

### Large changes in *N*_*e*_ detected in four species

Taking the results of the blockwise and PSMC analyses across all seven parasitoid species together, four species (*M. stigmatizans, M. dorsalis, O. nitidulus* and *T. auratus*) show evidence for large (at least a factor of three) changes in population size during the Quaternary period (Figure 1). In these species, an instantaneous step change in *N*_*e*_ fits the blockwise data significantly better than a null model of a constant *N*_*e*_. We infer a decrease in *N*_*e*_ towards the present (*λ <* 1) in three species (*M. stigmatizans, M. dorsalis* and *O. nitidulus*) and an increase in one species (*T. auratus*) (*λ >* 1) (Figure 1 and Table 1). In all four species, the change in *N*_*e*_ in the PSMC trajectory (Figure 1) agrees both in direction and timescale with the inference under the single step-change model. However, the *N*_*e*_ changes visible in the PSMC trajectory for these species are not equally abrupt. For example, *T. auratus* shows a gradual increase of *N*_*e*_ over a period of more than 200 ky, while the decreases in the PSMC trajectories of *M. dorsalis* and *M. stigmatizans* are comparatively sudden. With the exception of *M. stigmatizans*, the magnitude of *N*_*e*_ change inferred under the step-change model is greater than the relative magnitude of peak *N*_*e*_ in the PSMC trajectories (Figure 1, Tables 1 and 2). Inferences based on PSMC and blockwise analyses are also broadly similar for *O. pomaceus* and *C. fungosa*: a step change in *N*_*e*_ is supported in neither species, and both show comparatively small population size changes in the PSMC analysis (Tables 1 and 2).

**Figure 1:**
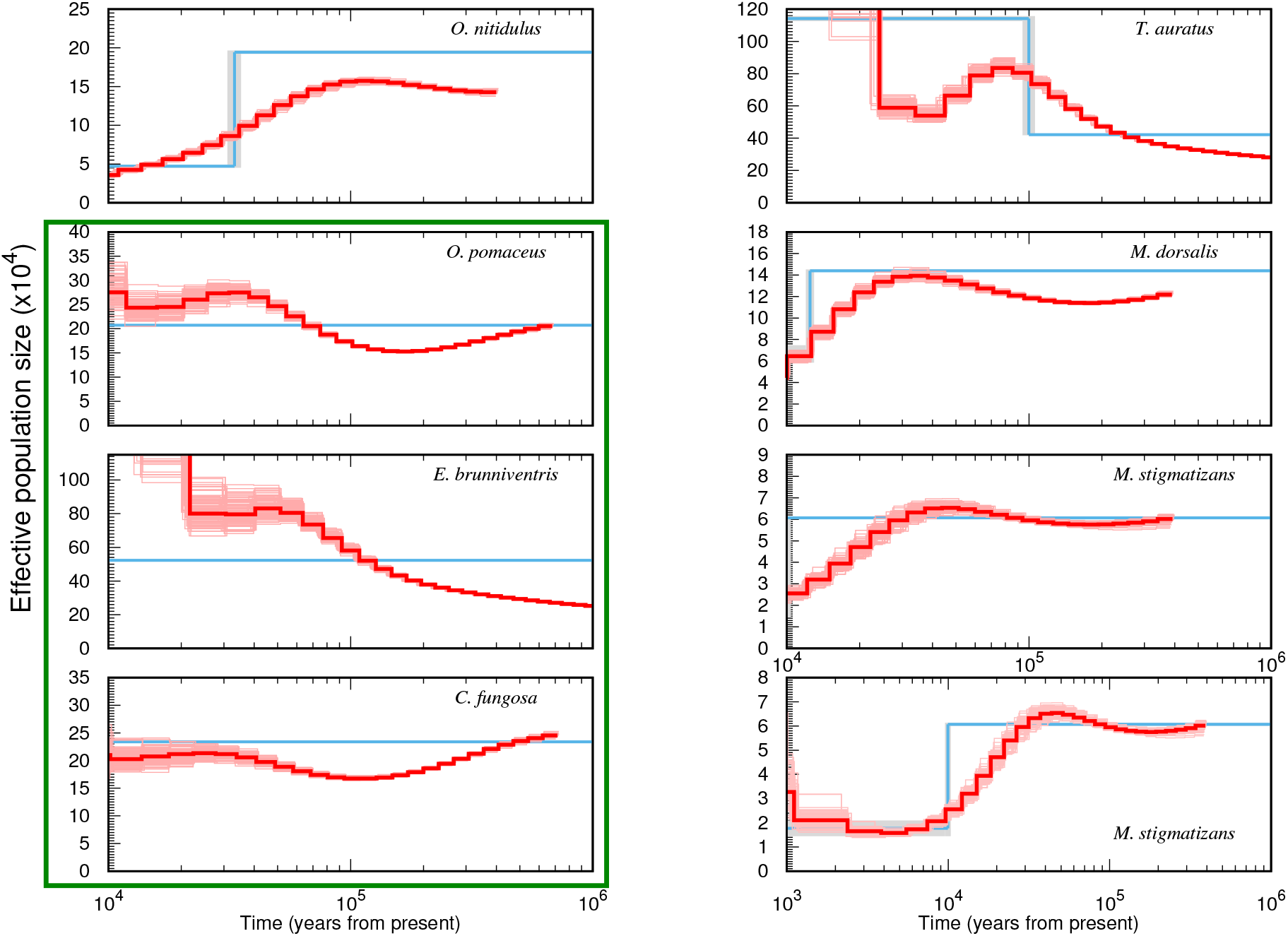
PSMC and maximum composite likelihood estimates (MCLE) of *N*_*e*_ for Iberian populations of seven species of chalcid parasitoid wasps. PSMC estimates and bootstrap replicates are shown in dark red and pale red, respectively. Population sizes estimated using the composite likelihood step change model are shown in blue with 95% CIs in grey. Species for which a step change model is not supported are outlined in green. Results for *M. stigmatizans* are also plotted on an alternative timescale to reveal recent *N*_*e*_ change (bottom right).

*E. brunniventris* is an exception to the broad agreement between blockwise and PSMC analyses: despite showing no support for a step change, the PMSC trajectory of this species indicates a substantial (but gradual) change in *N*_*e*_ (Table 2). However, additional analyses for this species (see Discussion) suggest that the PSMC analyses for *E. brunniventris* may be affected by genetic structure and/or the low contiguity of its reference genome.

### No support for temporal clustering of demographic events

The four species with support for a step change in *N*_*e*_ (*M. stigmatizans, M. dorsalis, O. nitidulus* and *T. auratus*) show no overlap in the estimated times of *N*_*e*_ change. Using an insect spontaneous mutation rate (Keightley et al., 2009) to calibrate *T* estimates (see Methods for caveats), Iberian populations of *M. stigmatizans* and *M. dorsalis* most likely decreased in size at the start of the current interglacial (10 and 12.5 kya respectively). In contrast, the decrease in *N*_*e*_ inferred for *O. nitidulus* most likely dates to late in the last glacial period (33 kya) and the increase in *N*_*e*_ in *T. auratus* at 100 kya dates to just after the end of the (Eemian) interglacial (Table 1).

Paralleling the blockwise model results, the times of peak *N*_*e*_ in the PSMC trajectories also had non-overlapping CIs (across bootstrap replicates) in *M. stigmatizans, M. dorsalis, O. nitidulus* and *T. auratus* (although values for the two *Megastigmus* species are close, see Table 1). Interestingly, the three species with no support for a step change (*O. pomaceus, C. fungosa* and *E. brunniventris*) as well *M. dorsalis* show highly similar (overlapping CI) times of peak *N*_*e*_ (Figure 2 and Table 2).

**Figure 2:**
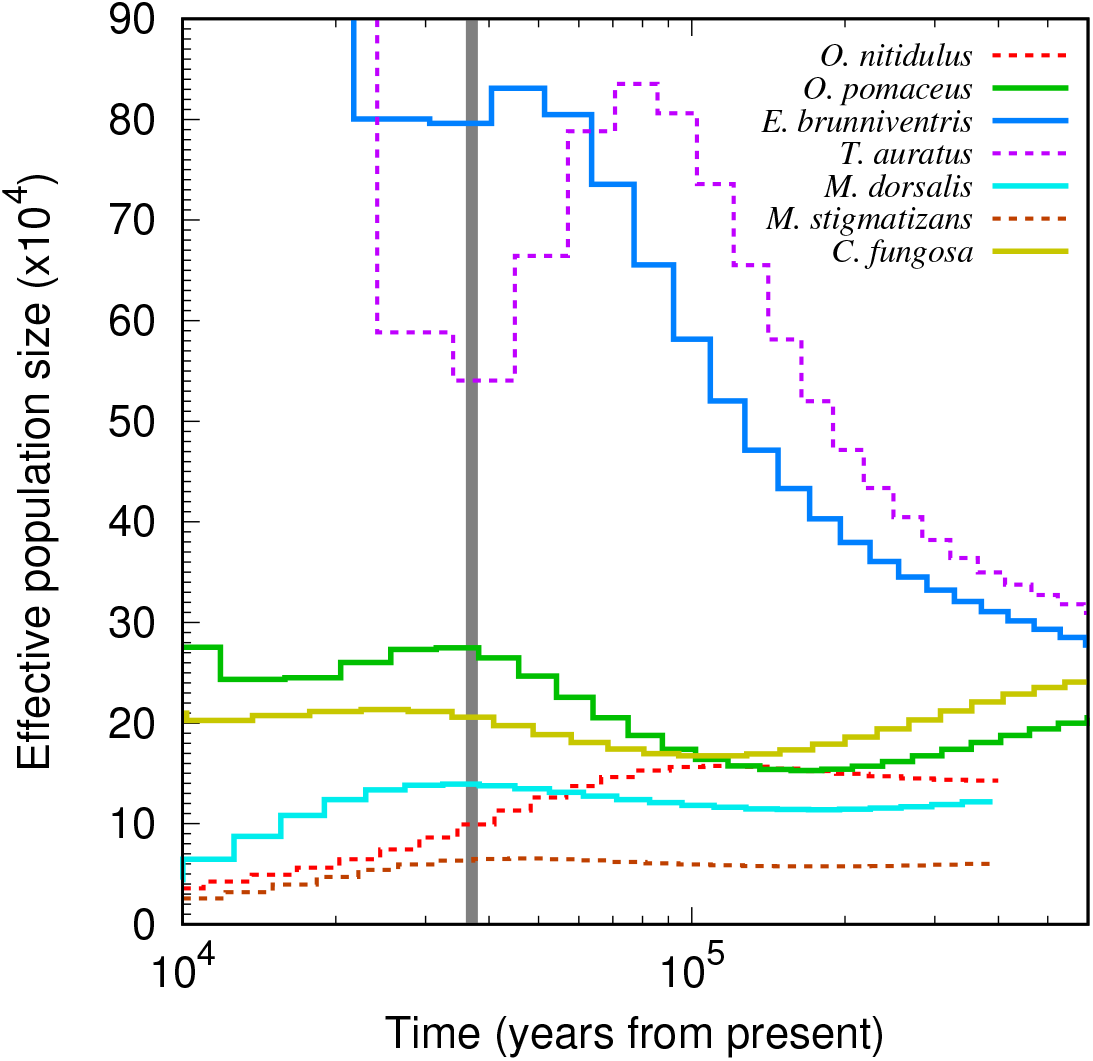
*N*_*e*_ trajectories inferred by PSMC for Iberian populations of seven species of parasitoids. The vertical grey bar indicates the overlap in peak *N*_*e*_ of bootstrap replicates across four species (solid lines) (*M. dorsalis, O. pomaceus, C. fungosa* and *E. brunniventris*). The remaining three species (dashed lines) have unique peak *N*_*e*_ times.

### Changes in *N*_*e*_ occur after the divergence of Iberian populations

In five species the PSMC trajectories of within-population (Iberia) pairs diverge clearly from the cross-population (Iberia *versus* Hungarian) trajectories. In contrast, little divergence between within and cross-population PSMC trajectories is visible for *E. brunniventris* and none for *T. auratus* (Figure 3). In the three species for which the blockwise analysis supports a decline in population size, the inferred time of *N*_*e*_ change is more recent than the divergence of within- and cross-population PSMC trajectories (Figure 3 and Table 2) and so must have occurred after Iberian and Balkan populations split. For *O. pomaceus*, *C. fungosa* and *M. stigmatizans* the split times between Iberian and the Balkan populations inferred here *post hoc* by comparing PSMCs trajectories are broadly compatible with the divergence estimates obtained by Bunnefeld et al. (2018) under explicit models of population divergence (Figure 3). In contrast, for both *M. dorsalis* and *O. nitidulus* PSMC based estimates of population divergence substantially predate the split times inferred by Bunnefeld et al. (2018). In both cases, the cross-population PSMC trajectories decrease after divergence between the Iberian and Hungarian populations, suggesting that these populations have been connected by some level of post-divergence gene flow.

**Figure 3:**
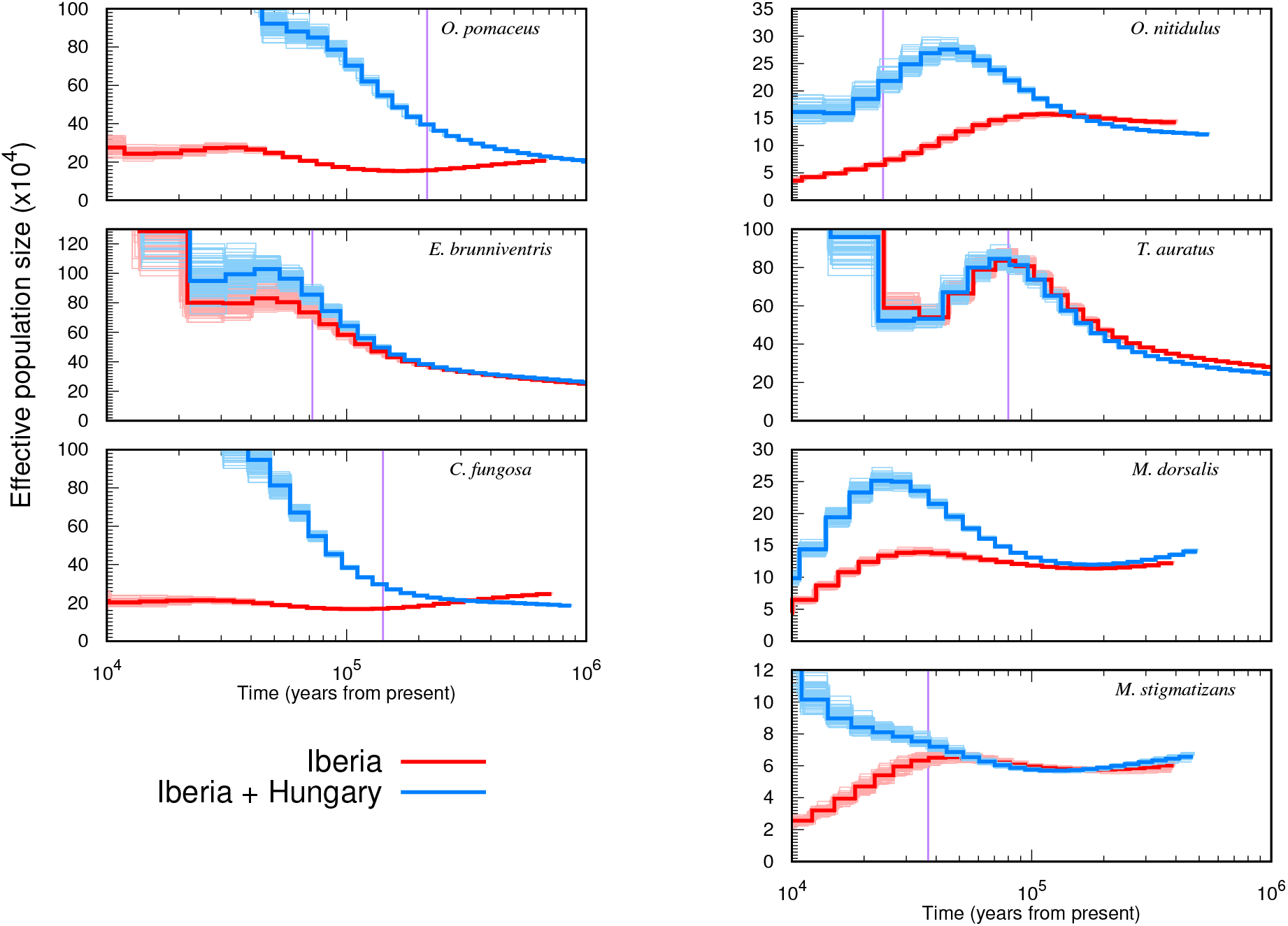
PSMC *N*_*e*_ trajectories for Iberian (red) and cross-population (Iberian vs Balkans) pairs (blue) of seven species of parasitoids. Population splits can be inferred as the time at which the trajectories of within-population and cross-population pairs diverge. Vertical bars indicate the split times estimated by Bunnefeld et al. (2018) using a MCL method based on the blockwise data. The split time estimated by Bunnefeld et al. (2018) for *M. dorsalis* is too recent (8 kya) to be visible in this figure.

### No evidence for structure within Iberia

We find little variation in PSMC trajectories between different Iberian pairs in almost all species (Figure S2), suggesting that our demographic inferences are unlikely to be substantially influenced by population structure within Iberia. For all species, with the exception of *E. brunniventris*, the differences in PSMC trajectories between Iberian pairs are similar in magnitude to the variation across bootstrap replicates and so likely reflect variation in coverage between individuals. *E. brunniventris* is the only species that showed clear variation in PSMC trajectories between Iberian pairs. While our additional analyses for *E. brunniventris* suggest other potential reasons for this finding (see Discussion), we cannot rule out that this variation is in part driven by population structure.

### Sensitivity analyses

Both demographic inference methods used here assume selective neutrality but make different simplifying assumptions about recombination: while PSMC approximates recombination as a Markov process along the genome, the blockwise composite likelihood framework assumes no recombination within blocks. To check the extent to which recombination and selection may bias parameter estimates, we fitted histories of a step change to data simulated with recombination. Our exploration of simulated data shows that both *N*_*e*_ and *λ* are underestimated with increasing recombination rate whereas the time of step change (*T*) is biased downwards when *N*_*e*_ declines towards the present (*λ <* 1) and upwards when *N*_*e*_ increases (*λ >* 1) (Table S3). These biases are an expected consequence of the shuffling of genealogical histories within a block in the presence of recombination, which reduces the variance in bSFS configurations. Given a recombination rate of *r* = 3 *×* 10^*−*9^ estimated for the parasitoids, our simulation check suggests that while estimates of *T*, the time of the step change may be underestimated by up to a factor of two, the ability to detect a step change and estimate its magnitude (*λ*) is little affected.

To investigate the potential effect of selective constraint on the block based inference we fitted a history of a single step change separately to genic and intergenic regions. Parameter estimates based on genic data for *O. nitidulus* were similar to the full data set (Table S2). While *N*_*e*_ is slightly lower and *T* is slightly greater for genic regions, as would be expected as a result of selective constraints, the timing and relative magnitude of demographic change was little affected by partitioning the data by annotation.

## Discussion

We analysed genome-wide sequence variation using two contrasting inference approaches to test if and how demographic histories vary within a guild of insect parasitoids in a single Pleistocene refugium. We find evidence for drastic declines in population size in three out of seven species (*M. stigmatizans, M. dorsalis, O. nitidulus*), and a large increase in population size one species (*T. auratus*). However, contrary to simple models of concordant community-wide responses to Pleistocene climatic events, we find that the demographic histories of species in this guild differ markedly both in direction and timescale.

Interestingly, the four species that show the smallest change in past population size overlap in terms of the time of peak *N*_*e*_. The apparent congruence in these species may well be due to a shared background demography, the signatures of which are masked in species that show more drastic and idiosyncratic changes in *N*_*e*_. However, given that PSMC assumes selective neutrality, which is problematic for compact insect genomes, the genome-wide effect of selection is an alternative (and not mutually exclusive) explanation for the shared timing of the “hump” in the PSMC trajectories. Schrider et al. (2016) have shown that selective sweeps can generate troughs in PSMC trajectories and background selection has been shown to lead to erroneous inference of population growth (Ewing and Jensen, 2016). We therefore cannot rule out the possibility that congruent peaks in *N*_*e*_ are an artefact of selective effects which, assuming similar genome composition, mutation, and recombination rates of these species, may distort PSMC inferences in similar ways. However, the fact that we have reconstructed a set of very different PSMC trajectories, most of which differ markedly from the selection-induced PSMC trajectories of Schrider et al. (2016) (in that they show large declines rather than increases in *N*_*e*_ toward the present), suggests that it is unlikely that selective effects are the main cause of the inferred *N*_*e*_ changes. For the blockwise analyses we find little difference in parameter estimates between genic and intergenic regions of *O. nitidulus* (Table S2), which again argues against a major effect of selection on our inferences. Furthermore, one would expect genomes with a shorter map length (physical length *×* recombination rate) to be disproportionately affected by linked selection (Smith and Haigh, 1974; Mackintosh et al., 2019). However, if anything we observe the opposite pattern: the two species with shortest map lengths (*O. nitidulus* and *C. fungosa*) show less pronounced *N*_*e*_ change than the two species with the longest map length (*M. stigmatizans* and *M. dorsalis*).

### Concordance between step change and PSMC analyses

PSMC and blockwise analyses exploit different aspects of the data and differ drastically in sampling schemes (contiguous pairwise alignments vs short blocks sampled across five individuals) and underlying assumptions: while the blockwise composite likelihood framework fits a single instantaneous step change in *N*_*e*_, PSMC reconstructs population size change as a continuous trajectory. Despite these differences, both methods yield broadly congruent conclusions: the four species for which the blockwise analyses diagnose an abrupt change in *N*_*e*_ also show PSMC trajectories with large *N*_*e*_ changes in the same direction and at similar times. The slightly greater magnitude of *N*_*e*_ change under the step change history compared to the PSMC analyses may be a consequence of the fact that larger samples contain more information about recent demography. In other words, the change in *N*_*e*_ in the recent past detectable with larger samples (five haploid individuals) in the blockwise framework may simply not be detectable by PSMC analysis (two individuals).

In general, one may view the fact that PSMC is essentially assumption-free as an advantage over model based inference, because it enables straightforward inference of *N*_*e*_ changes. Comparison of PSMC trajectories between pairs of individuals and populations allows qualitative assessment of divergence and admixture (Figure 3). However, the flip-side of this flexibility is that PSMC has limited resolution in the recent past and provides no obvious route for quantitatively testing (necessarily) simple demographic hypotheses across species. We have sought to overcome the latter limitation by comparing PSMC trajectories in terms of simple summaries (such as the timing and relative magnitude of peak *N*_*e*_).

### Changes in population size coincide with late Pleistocene climatic transitions

Our estimates of both the timing of drastic *N*_*e*_ changes and the cluster of peak *N*_*e*_ coincide with climatic events in the Quaternary: the start of the Holocene around 11 kya, a Dansgaard-Oeschger interstadial event – a brief warm period during the last glacial – 37 kya and the end of the Eemian interglacial around 106 kya. Previous studies on these and other species in the gallwasp community have inferred an increase in gene flow between refugia during these three time periods (Bunnefeld et al., 2018; Lohse et al., 2010, 2012). In particular, the cluster of peak *N*_*e*_ times during the Dansgaard-Oeschger interstadial event coincides with a large community-wide peak in the frequency of admixture between refugia inferred by Bunnefeld et al. (2018). Similarly, the increase in *N*_*e*_ for *T. auratus* at the end of the Eemian interglacial period coincides with the divergence of the Iberian population of this species inferred previously (Bunnefeld et al., 2018; Stone et al., 2012). It seems plausible that both events are associated with the geographic expansion of suitable oak habitat across refugial barriers during these times (Brewer et al., 2002; Petit et al., 2002), which may have triggered parasitoid range expansions.

### Population structure within Iberia and gene flow from Eastern refugia

Population structure within southern European refugia has previously been demonstrated for some species, and it has been suggested that given its complex topography Iberia should be considered a mosaic of multiple micro-refugia rather than a single entity (Feliner, 2011; Hearn et al., 2014). However, our PSMC results suggest a complete lack of population structure in six out of seven species in the parasitoid guild (Figure S2) and imply high gene flow across Iberia. This is perhaps unsurprising given that gallwasp-associated parasitoid wasps (and other chalcids) are able to disperse long distances even across patchy habitats and host distributions (Hayward and Stone, 2006; Compton et al., 2000).

Our estimates of population split times based on comparisons of within and cross-population PSMC trajectories agree broadly with those of Bunnefeld et al. (2018) for most species. *M. dorsalis* and *O. nitidulus*, the two species that show the least agreement with past estimates, both show decreases in cross-population *N*_*e*_ after divergence that are compatible with ongoing gene flow between refugia. The model space considered by Bunnefeld et al. (2018) was limited to histories involving a single burst of instantaneous admixture between refugial populations, with no potential to detect such post divergence gene flow.

### *E. brunniventris* is an outlier

*E. brunniventris* is an outlier in our results in several ways: it is the species with the highest genetic diversity, shows signals of population structure and is the only species for which our two inference approaches disagree. While the blockwise analysis gives no support for a change in *N*_*e*_, the PSMC trajectory shows a steady increase by a factor of four. To explore the ability of the blockwise analyses to detect gradual changes in population size, we simulated 100 replicate data sets (assuming the same size and block length as the real data) under the gradual change in *N*_*e*_ inferred via PSMC for *E. brunniventris*. We find that in each case, a history of a single step change fits the data better than a null model of constant *N*_*e*_, indicating that the blockwise analysis is indeed sensitive to gradual increases in population size. A possible explanation for the discrepancy between the PSMC and blockwise inferences for *E. brunniventris* is that its genome assembly, the least contiguous among our set of taxa (N50 of 3.8kb, Table S1), is simply too fragmented for reliable PSMC inference. Since by default PSMC does not consider contigs *<* 10kb long, PSMC inference for *E. brunniventris* is based on a likely conserved subset of the genome. However, even when the blockwise analysis is limited to contigs *>* 10kb, a null model of constant *N*_*e*_ cannot be rejected. Interestingly, both a history of constant *N*_*e*_ and a step change model give a poor fit to the observed frequency of bSFS configurations in *E. brunniventris*. In particular, *E. brunniventris* shows an excess of both monomorphic blocks and blocks with a large number of variants (Figure S4), suggesting that its history is not well approximated by any model that assumes a single panmictic population. The lack of divergence of within and cross-population PSMC trajectories for *E. brunniventris* would be compatible with substantial gene flow between the Iberian and Hungarian populations (Figure 3). Alternatively *E. brunniventris* – an extreme generalist attacking a wide range of oak gallwasp hosts (Askew et al., 2013) – may harbour genetic structure as a result of recent divergence into cryptic host races. However, in the absence of a better reference genome and larger samples it remains unclear to what extent the disagreement between PSMC and blockwise analyses for *E. brunniventris* is indicative of a more complex history.

### Outlook

We have uncovered considerable diversity in the demographic histories of a guild of insect parasitoids at the scale of a single glacial refugium. Our results mirror the findings of Bunnefeld et al. (2018) who showed that the relationships between refugia in this guild are to a large extent idiosyncratic. An obvious question is to what extent differences in demographic history correlate with ecological traits. Intriguingly, the four parasitoid species with support for a step change (in either direction) have a narrower host range (Askew et al., 2013) and a lower ancestral *N*_*e*_ than those for which a null model of constant *N*_*e*_ could not be rejected (Figure S3, Table S1). Although these trends are not significant, they are compatible with the idea that specialists are more prone to large changes in *N*_*e*_ than generalists and, as a consequence, may also be at a higher risk of extinction (Colles et al., 2009; McKinney, 1997). Larger samples of species, incorporating wide diversity in host number and other relevant traits (such as dispersal ability) are required to explore the potential relationships between species characteristics and demographic history.

Parasitoid wasps are particularly suited for such systematic comparisons across co-distributed species given their rich biology and manageable genomes, which can be sampled in a haploid state by targeting males. It will be particularly interesting to interrogate such data with a new generation of inference approaches that reconstruct ancestral recombination graphs (ARG) from phased genomes (Kelleher et al., 2019; Speidel et al., 2019). Given sufficiently large samples, it may be possible to infer much more detailed trajectories of *N*_*e*_ change and population divergence times directly from such reconstructed ARGs.

## Acknowledgements

We thank Lynsey Bunnefeld for help in the molecular lab and useful discussions and José Luis Nieves-Aldrey and Juli Pujade-villar for contributing samples and Sam Ebdon for comments on the manuscript. This work was supported by a Natural Environment Research Council to GNS and KL (NE/J010499/1). WW was supported by an E3 doctoral training studentship from the Natural Environment Research Council (NERC) UK, KL is supported by a NERC fellowship (NE/L011522/1) and an ERC starting grant (ModelGenomLand).

## Data Accessibility

- Raw reads have been deposited in the European Nucleotide Archive (ENA) (ERP023079) and the SRA (PRJEB20883)
- Genome assemblies are deposited in the ENA (PRJEB27189 and ERP109243)
- *Mathematica* notebook and blockwise data are available as Supporting Information

## Author Contributions

KL and GS designed the project; WW analysed the sequence data with contributions from KL; all authors wrote the manuscript.

## Supplementary information

**Figure S1:**
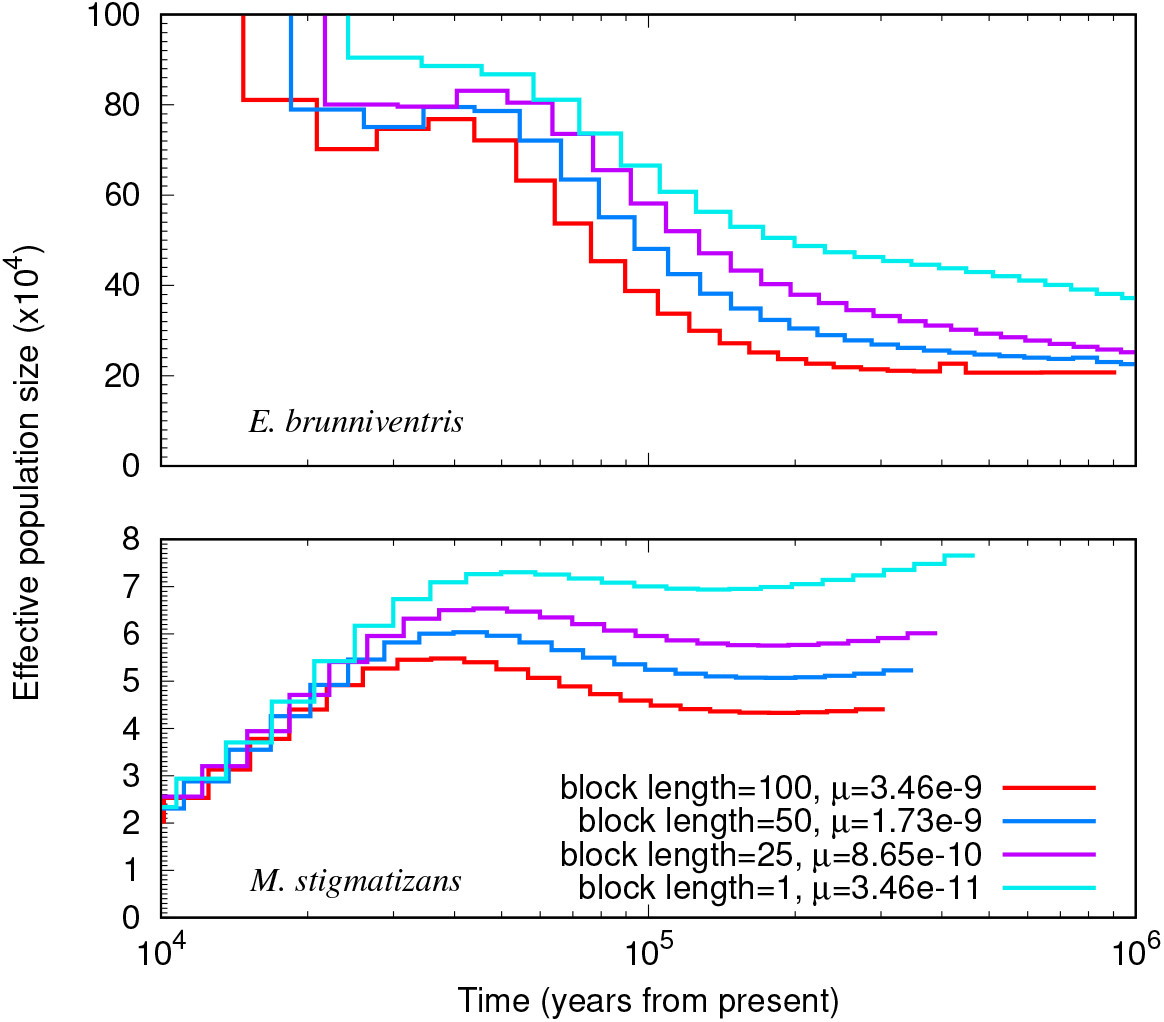
*N*_*e*_ trajectories inferred by PSMC using different block lengths for *E. brunniventris* and *M. stigmatizans*. The mutation rate is adjusted accordingly for each block length.

**Figure S2:**
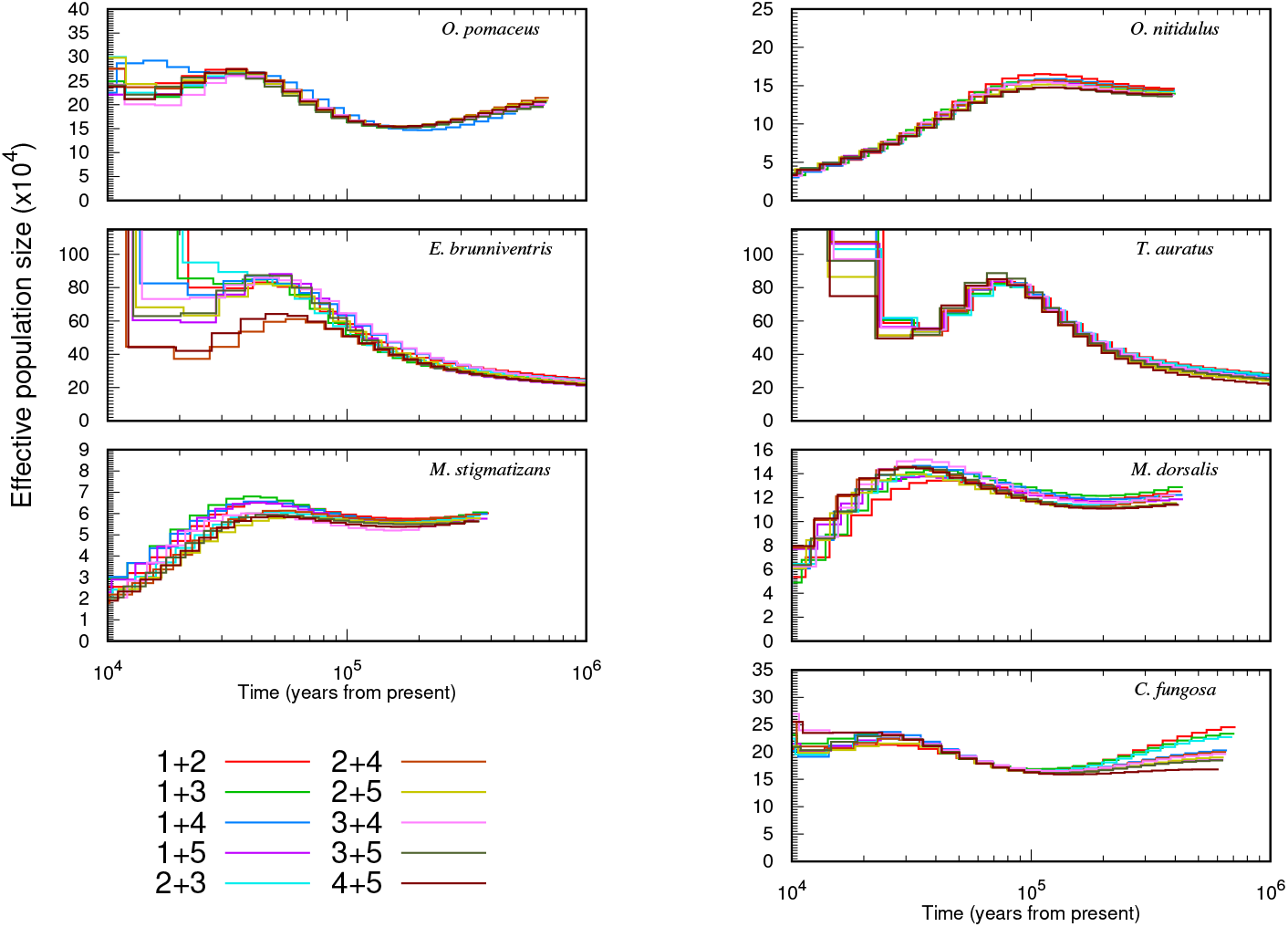
*N*_*e*_ trajectories inferred by PSMC for all pairwise combinations of the five Iberian haploid males of each species. Individuals 1 and 2 for each species correspond to the focal pair in figures 1 2.

**Table S1:**
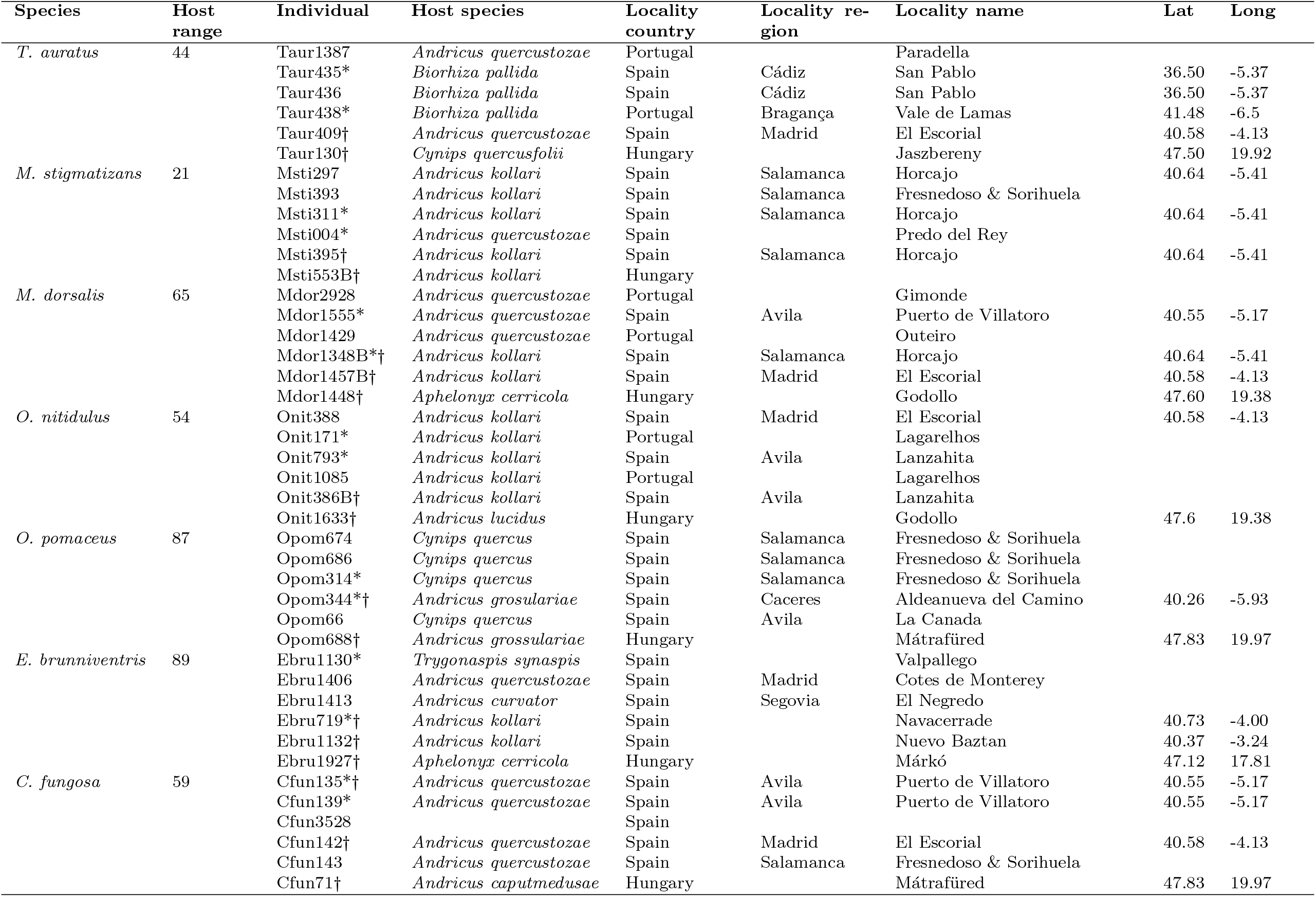
Sampling locations and gallwasp hosts of parasitoid individuals used for whole genome sequencing. *: The two focal Iberian individuals used for PSMC analysis *†*: Individuals previously analysed by Bunnefeld et al. (2018)

**Table S2:** Simulation investigation of recombination bias in the likelihood method. Recombination rate for simulations with varying *λ* and block length is 2.3 × 10^*−*9^. Comparison of genic and intergenic regions is performed on real data

| Independent parameter | $\theta$ | $\lambda$ | $T$ |
| --- | --- | --- | --- |
| recombination rate (/bp/generation) |  |  |  |
| input parameters: | 1.5 | 0.5 | 0.5 |
| 0 | 1.509 | 0.480 | 0.492 |
| $3.46 \times 10^{-11}$ | 1.520 | 0.498 | 0.500 |
| $2.3 \times 10^{-10}$ | 1.518 | 0.502 | 0.470 |
| $7.3 \times 10^{-10}$ | 1.525 | 0.506 | 0.425 |
| $2.3 \times 10^{-9}$ | 1.457 | 0.497 | 0.281 |
| $7.3 \times 10^{-9}$ | 1.195 | 0.420 | 0.100 |
| $2.3 \times 10^{-8}$ | 1.885 | 0.668 | 0.007 |
| $\lambda$ | | | |
| input parameters: | 1.0 |  | 1.0 |
| 0.1 | 1.130 | 0.084 | 0.864 |
| 0.25 | 1.089 | 0.292 | 0.809 |
| 0.5 | 1.052 | 0.622 | 0.662 |
| 0.75 | 0.912 | 0.794 | 0.100 |
| 1 | 1.067 | 1.307 | 1.275 |
| 1.25 | 1.049 | 1.617 | 1.165 |
| 1.5 | 1.046 | 1.909 | 1.067 |
| 2 | 1.032 | 2.636 | 1.077 |
| 2.5 | 1.025 | 3.289 | 1.055 |
| genomic regions of *O. nitidulus* |  |  |  |
| whole genome | 0.381 | 0.245 | 0.683 |
| intergenic | 0.401 | 0.265 | 0.591 |
| genic | 0.359 | 0.231 | 0.903 |

**Table S3:**
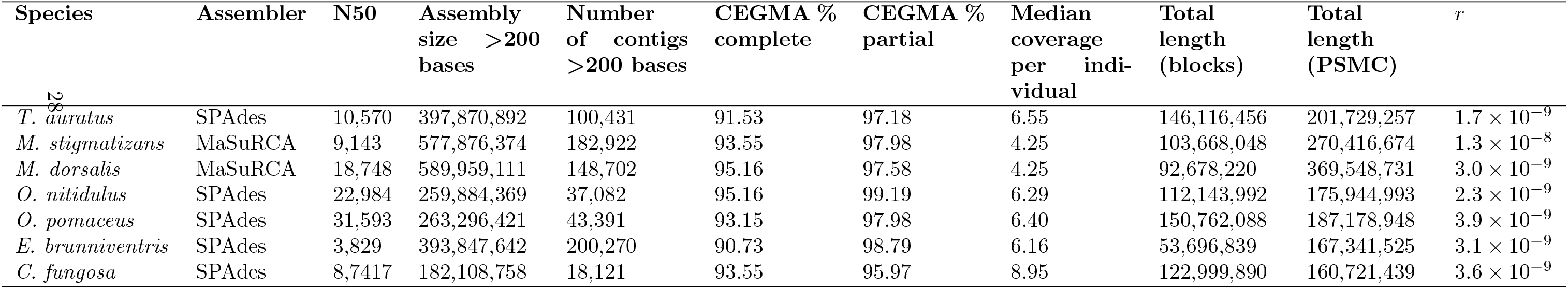
Assembly summaries. N50: 50% of the assembly is contained in contigs of length equal to or greater than this value. CEGMA (Core Eukaryotic Genes Mapping Approach): full or partial presence of a core set of eukaryotic genes. *r*: recombination rate per base pair per generation estimated by Bunnefeld et al. (2018).

**Figure S3:**
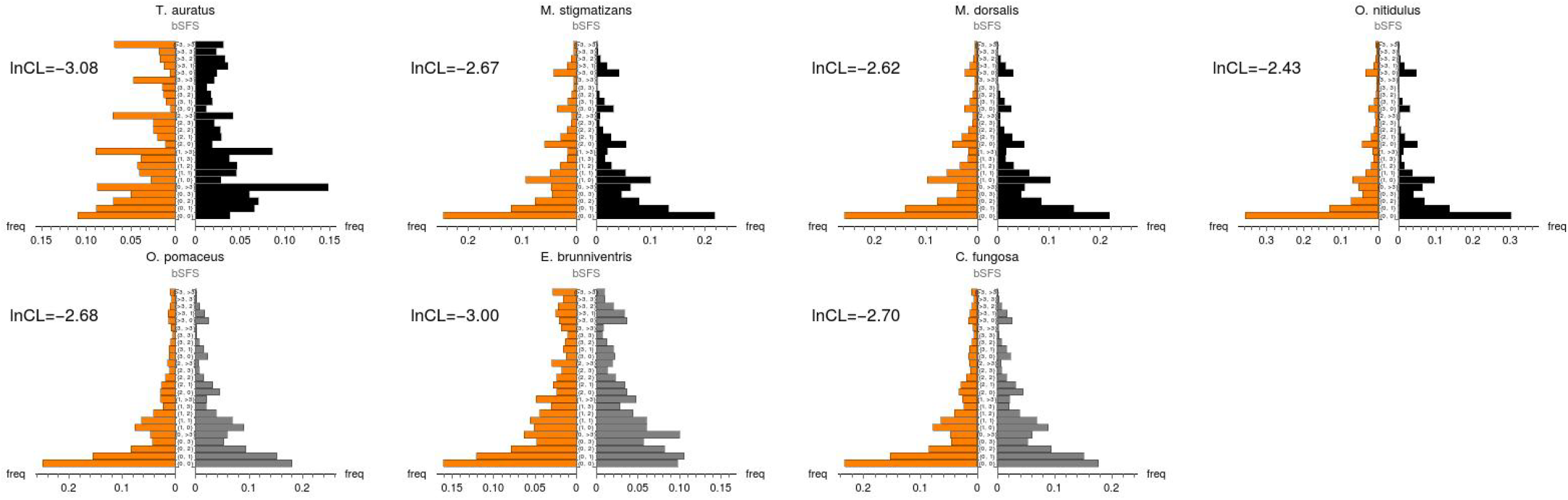
Goodness of fit of the best fitting demographic histories for seven species of parasitoids. The observed frequencies of blockwise configurations defined via the folded SFS are shown in orange (left). The expected frequency under the best fitting history is shown on the right for four species fitting a step change model (top, black) and three species for which a constant *N*_*e*_ could not be rejected (bottom, gray). For a sample of *n* = 5 the folded SFS has two entries *j*_2,3_, *j*_1,4_. The log composite likelihood, *lnCL* (per block), a measure of absolute goodness of fit of the data to the inferred model, is shown for each species.

**Figure S4:**
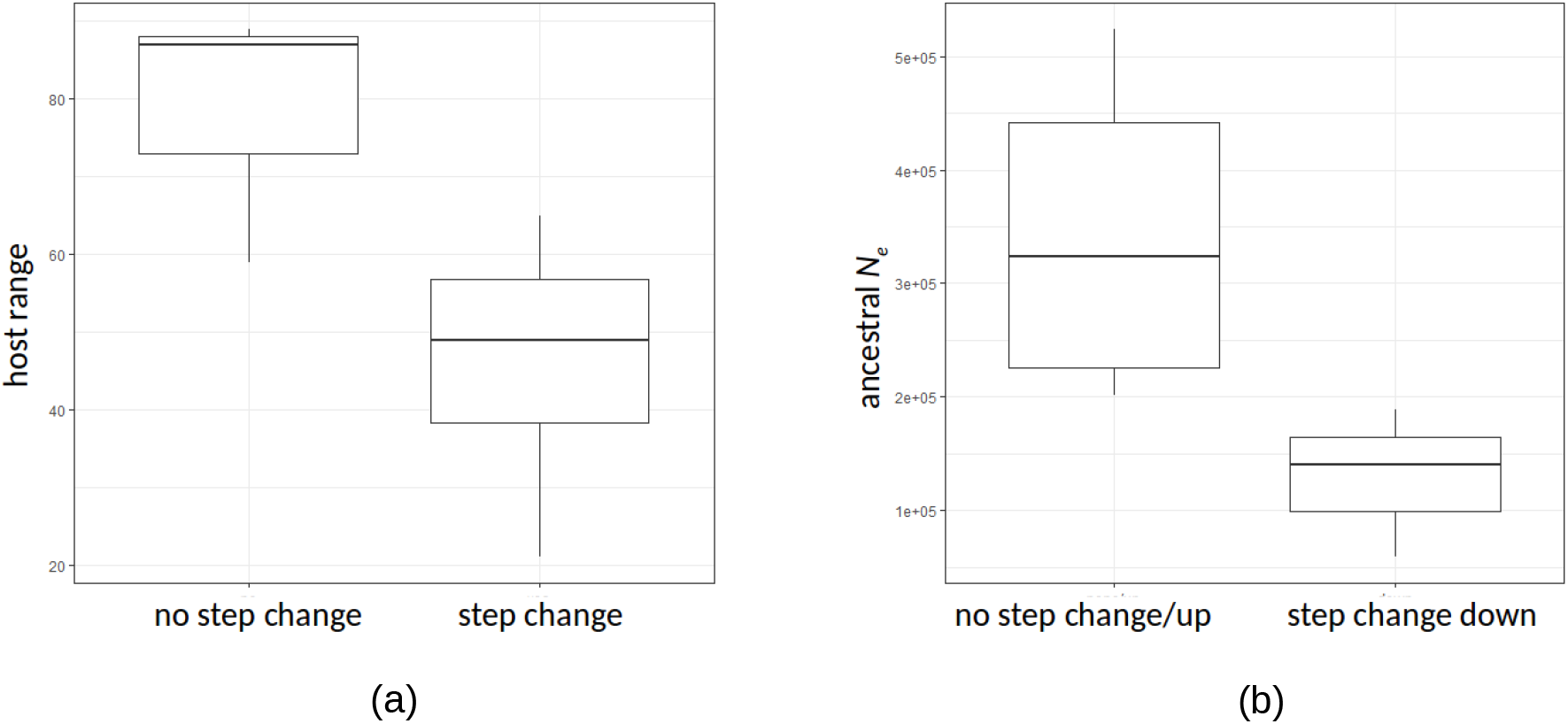
(a) Host range and (b) ancestral *N*_*e*_ of species with and without *N*_*e*_ changes inferred by the likelihood method.

