## Supporting Material 1 for "Diverse demographic histories in a guild of hymenopteran parasitoids"

### Deriving the GF under a step Change model, n=5

We use the automation for the generating funtion (GF) implemented by Lohse et al. 2016 (Supporting notebook S1) to derive the generating function for a history of a single step change:

In[167]:=
```
Step5GFsol = MakeSolvedGF[ψ[ω, {{a}, {b}, {c}, {d}, {e}},
    ChangeCoalescence → {{d1, λ1}}], {{X}, {X}, {X}, {X}, {X}}];
```

In[168]:=
```
Step5GFsol /. ω[_] → 0 // Simplify
```

Out[168]= $\left\{\frac{1}{3}, \frac{1}{6}, \frac{1}{2}\right\}$

In[169]:=
```
Step5GFsolE = (unRoot[Step5GFsol, ω] /. EquivalenceRules[
      {{a}, {b}, {c}, {d}, {e}}, {{X}, {X}, {X}, {X}, {X}}]) // Simplify;
```

Inverting wrt to the time of the step change:

In[170]:=
```
Step5GFsolEInv =
  (((InverseLaplaceTransform[(d1[]) ^-1 * #, d1[], T1]) & /@ Step5GFsolE) /.
     λ1[] → λ1) // Simplify;
```

In[171]:=
```
Step5GFsolEInv /. ω[_] → 0
```

Out[171]= $\left\{\left\{\frac{1}{3}, \frac{1}{6}, \frac{1}{2}\right\}\right\}$

Saving inverted GF and assigning it to an object:

In[172]:=
```
Step5GFsolEInv >>
 "/home/konrad/Dropbox/Manuscripts/Walton_Iberian_demography_2019/StepUnroot5.m"
```

In[173]:=
```
testtab = likFullTab[Step5GFsolEInv // Total, ω, 1.1, {T1 → 0.2, λ1 → 0.5}, 3]
```

Out[173]= SparseArray[ Specified elements: 25 Dimensions: {5, 5} ]

In[174]:=
```
testtab // Normal // TableForm
```

Out[174]//TableForm=

| | | | | |
|---|---|---|---|---|
| 0.0846039 | 0.0534902 | 0.0362508 | 0.025109 | 0.0560377 |
| 0.0820987 | 0.0487533 | 0.0309913 | 0.0209014 | 0.046407 |
| 0.0651872 | 0.036427 | 0.021136 | 0.0134486 | 0.0289007 |
| 0.0485377 | 0.0260755 | 0.0138342 | 0.00811548 | 0.0162502 |
| 0.115534 | 0.0603876 | 0.0282109 | 0.0136721 | 0.0196393 |

In[175]:=
```
testtab // Normal // Flatten // Total
```

Out[175]= 1.

### Importing and summarising data

Load parasitoid block data

In[192]:=
```
taur = Get[
   "/home/konrad/Dropbox/Manuscripts/Walton_Iberian_demography_2019/blockwise_data
     _files/Taur_nton_counts_table_KL_full_2.txt"];
msti = Get[
   "/home/konrad/Dropbox/Manuscripts/Walton_Iberian_demography_2019/blockwise_data
     _files/Msti_nton_counts_table_KL_full_2.txt"];
mdor = Get[
   "/home/konrad/Dropbox/Manuscripts/Walton_Iberian_demography_2019/blockwise_data
     _files/Mdor_nton_counts_table_KL_full_2.txt"];
onit = Get[
   "/home/konrad/Dropbox/Manuscripts/Walton_Iberian_demography_2019/blockwise_data
     _files/Onit_nton_counts_table_KL_full_2.txt"];
opom = Get[
   "/home/konrad/Dropbox/Manuscripts/Walton_Iberian_demography_2019/blockwise_data
     _files/Opom_nton_counts_table_KL_full_2.txt"];
ebru = Get[
   "/home/konrad/Dropbox/Manuscripts/Walton_Iberian_demography_2019/blockwise_data
     _files/Ebru_nton_counts_table_KL_full_2.txt"];
cfun = Get[
   "/home/konrad/Dropbox/Manuscripts/Walton_Iberian_demography_2019/blockwise_data
     _files/Cfun_nton_counts_table_KL_full_2.txt"];
```

In[199]:=
```
allSp = {"T. auratus", "M. stigmatizans", "M. dorsalis",
   "O. nitidulus", "O. pomaceus", "E. brunniventris", "C. fungosa"};
```

In[200]:=
```
allblocks = {taur, msti, mdor, onit, opom, ebru, cfun};
allblockl = {328, 1584, 558, 586, 376, 223, 338};
```

The total length of sequence (bases) and pairwise $\pi$:

In[204]:=
```
{allSp, ll = ((Length[#] & /@ allblocks) * allblockl),
  piall = ((pi[#] & /@ allblocks) / ll)} // TableForm
```

Out[204]//TableForm=

| | | | | | |
|---|---|---|---|---|---|
| T. auratus | M. stigmatizans | M. dorsalis | O. nitidulus | O. pomaceus | E. bru |
| 146 116 456 | 103 668 048 | 92 678 220 | 112 143 992 | 150 762 088 | 53 696 [illegible] |
| 0.0050907 | 0.000540988 | 0.0014261 | 0.0012645 | 0.00230131 | 0.0056[illegible] |

The expected fSFS for a population of constant $N_e$ is:

In[205]:=
```
sfsExpt = Total[Flatten[-D[Step5GFsolEInv /. {λ1 → 1, T1 → 0}, #] /. ω[_] → 0]] & /@
    Drop[Variables[Step5GFsolEInv], 1];
```

In[206]:=
```
sfsExpt / Total[sfsExpt] // N
```

Out[206]= {0.6, 0.4}

Three out of four species have an excess of intermediate frequency variants:

In[207]:=
```
TableForm[{allSp, SetPrecision[sfsnorm[#] & /@ allblocks, 3]}, TableDepth → 2]
```

Out[207]//TableForm=

| | | | | |
|---|---|---|---|---|
| T. auratus | M. stigmatizans | M. dorsalis | O. nitidulus | O. pomaceus |
| {0.677, 0.323} | {0.525, 0.475} | {0.560, 0.440} | {0.575, 0.425} | {0.627, 0.373} |

### Summarizing blockwise configurations

In[208]:=
```
allcounts = configCounts[#, 3] & /@ allblocks;
```

### Fitting a null model of constant $N_e$:

Fitting a null model:

In[•]:=
```
testnull =
  NMaximize[{totallnL[Step5GFsolEInv // Total, ω, theta, {λ1 → 1, T1 → 0}, 3, #],
      3 > theta > 0.5}, {theta}, MaxIterations → 150,
     Method → {"NelderMead", "ShrinkRatio" → 0.5, "ContractRatio" → 0.5,
       "ReflectRatio" → 1, "PostProcess" → False}] & /@ allcounts
```

Out[•]=
```
{{-1.37491 × 10^6, {theta → 2.36927}},
 {-176 241., {theta → 1.0297}}, {-436 000., {theta → 0.969765}},
 {-474 719., {theta → 0.867027}}, {-1.07508 × 10^6, {theta → 1.0788}},
 {-721 353., {theta → 1.61345}}, {-983 807., {theta → 1.09451}}}
```

In all cases, pairwise $\pi$ is lower than the estimate of $\theta$:

In[•]:=
```
{allSp, (#[[2, 1, 2]] & /@ testnull) / allblockl, ((pi[#] & /@ allblocks) / ll)} // TableForm
```

Out[•]//TableForm=

| | | | | | | |
|---|---|---|---|---|---|---|
| Taur | Msti | Mdor | Onit | Opom | Ebru | ( |
| 0.00722338 | 0.000650065 | 0.00173793 | 0.00147957 | 0.00286914 | 0.00723521 | ( |
| 0.0050907 | 0.000540988 | 0.0014261 | 0.0012645 | 0.00230131 | 0.00568784 | ( |

### Fitting a single step change in $N_e$

A model with three parameters:

In[209]:=
```
AbsoluteTiming[taurStep =
   NMaximize[{totallnL[Step5GFsolEInv // Total, ω, theta, {}, 3, allcounts[[1]]],
     5.9 > theta > 0.1 && 4.5 > λ1 > 0 && 2.0 > T1 > 0}, {theta, λ1, T1},
    MaxIterations → 150, Method → {"NelderMead", "ShrinkRatio" → 0.5,
      "ContractRatio" → 0.5, "ReflectRatio" → 1, "PostProcess" → False}]]
```

Out[209]=
```
{282.348, {-1.37004 × 10^6, {theta → 5.18393, λ1 → 2.70729, T1 → 0.0875168}}}
```

```
In[210]:= AbsoluteTiming[mstiStep =
    NMaximize[{totallnL[Step5GFsolEInv // Total, ω, theta, {}, 3, allcounts[[2]]],
      3 > theta > 0.001 && 3 > λ1 > 0.001 && 3 > T1 > 0}, {theta, λ1, T1},
     MaxIterations → 150, Method → {"NelderMead", "ShrinkRatio" → 0.5,
       "ContractRatio" → 0.5, "ReflectRatio" → 1, "PostProcess" → False}]]

    NMaximize: Failed to converge to the requested accuracy or precision within 150 iterations.

Out[210]= {285.075, {-174 688., {theta → 0.388485, λ1 → 0.292086, T1 → 0.281174}}}

In[211]:= AbsoluteTiming[mdorStep =
    NMaximize[{totallnL[Step5GFsolEInv // Total, ω, theta, {}, 3, allcounts[[3]]],
      3 > theta > 0.01 && 3 > λ1 > 0.01 && 4.0 > T1 > 0}, {theta, λ1, T1},
     MaxIterations → 150, Method → {"NelderMead", "ShrinkRatio" → 0.5,
       "ContractRatio" → 0.5, "ReflectRatio" → 1, "PostProcess" → False}]]

    NMaximize: Failed to converge to the requested accuracy or precision within 150 iterations.

Out[211]= {284.405, {-435 049., {theta → 0.514239, λ1 → 0.462565, T1 → 0.18722}}}

In[212]:= AbsoluteTiming[onitStep =
    NMaximize[{totallnL[Step5GFsolEInv // Total, ω, theta, {}, 3, allcounts[[4]]],
      2.5 > theta > 0.01 && 2.5 > λ1 > 0 && 2.0 > T1 > 0}, {theta, λ1, T1},
     MaxIterations → 150, Method → {"NelderMead", "ShrinkRatio" → 0.5,
       "ContractRatio" → 0.5, "ReflectRatio" → 1, "PostProcess" → False}]]

Out[212]= {283.34, {-464 762., {theta → 0.382493, λ1 → 0.242695, T1 → 0.702854}}}

In[213]:= AbsoluteTiming[opomStep =
    NMaximize[{totallnL[Step5GFsolEInv // Total, ω, theta, {}, 3, allcounts[[5]]],
      2.9 > theta > 0.1 && 2.9 > λ1 > 0 && 10 > T1 > 0.1}, {theta, λ1, T1},
     MaxIterations → 150, Method → {"NelderMead", "ShrinkRatio" → 0.5,
       "ContractRatio" → 0.5, "ReflectRatio" → 1, "PostProcess" → False}]]

Out[213]= {84.7199, {-1.07508 × 10^6, {theta → 1.07876, λ1 → 1.02711, T1 → 8.69084}}}

In[214]:= AbsoluteTiming[ebruStep =
    NMaximize[{totallnL[Step5GFsolEInv // Total, ω, theta, {}, 3, allcounts[[6]]],
      3.9 > theta > 0.01 && 2.9 > λ1 > 0.01 && 4 > T1 > 0}, {theta, λ1, T1},
     MaxIterations → 150, Method → {"NelderMead", "ShrinkRatio" → 0.5,
       "ContractRatio" → 0.5, "ReflectRatio" → 1, "PostProcess" → False}]]

Out[214]= {211.436, {-721 197., {theta → 1.59012, λ1 → 0.013708, T1 → 3.04265}}}

In[215]:= AbsoluteTiming[taurStep =
    NMaximize[{totallnL[Step5GFsolEInv // Total, ω, theta, {}, 3, allcounts[[7]]],
      3 > theta > 0.01 && 2 > λ1 > 0.01 && 3.5 > T1 > 0}, {theta, λ1, T1},
     MaxIterations → 150, Method → {"NelderMead", "ShrinkRatio" → 0.5,
       "ContractRatio" → 0.5, "ReflectRatio" → 1, "PostProcess" → False}]]

Out[215]= {194.087, {-983 688., {theta → 1.08286, λ1 → 0.303694, T1 → 3.49994}}}

In[216]:= allStep = {taurStep, mstiStep, mdorStep, onitStep, opomStep, ebruStep, taurStep};
```

### The residual

We can compare the observed bSFS with the expectation under the best fitting history:

In[∘]:=
```
expbSFS = Table[likFullTab[Step5GFsolEInv // Total,
    ω, allStep⟦i, 2, 1, 2⟧, Drop[allStep⟦i, 2⟧, 1], 3], {i, 1, 7}];
expbSFSflat = Table[expbSFS⟦i⟧ // Normal // Thread // Flatten, {i, 1, 7}];
obsbSFSflat = Table[(allcounts⟦i⟧ // Normal // Thread // Flatten) /
     Total[(allcounts⟦i⟧ // Normal // Flatten)] // N, {i, 1, 7}];
```

In[∘]:=
```
GraphicsGrid[{Table[ListPlot[{expbSFSflat⟦i⟧, obsbSFSflat⟦i⟧} // Thread,
    AxesLabel → {"exp", "obs"}, PlotLabel → allSp⟦i⟧], {i, 1, 4}],
  Table[ListPlot[{expbSFSflat⟦i⟧, obsbSFSflat⟦i⟧} // Thread,
    AxesLabel → {"exp", "obs"}, PlotLabel → allSp⟦i⟧], {i, 5, 7}]}, PlotRange → All]
```

Out[∘]=

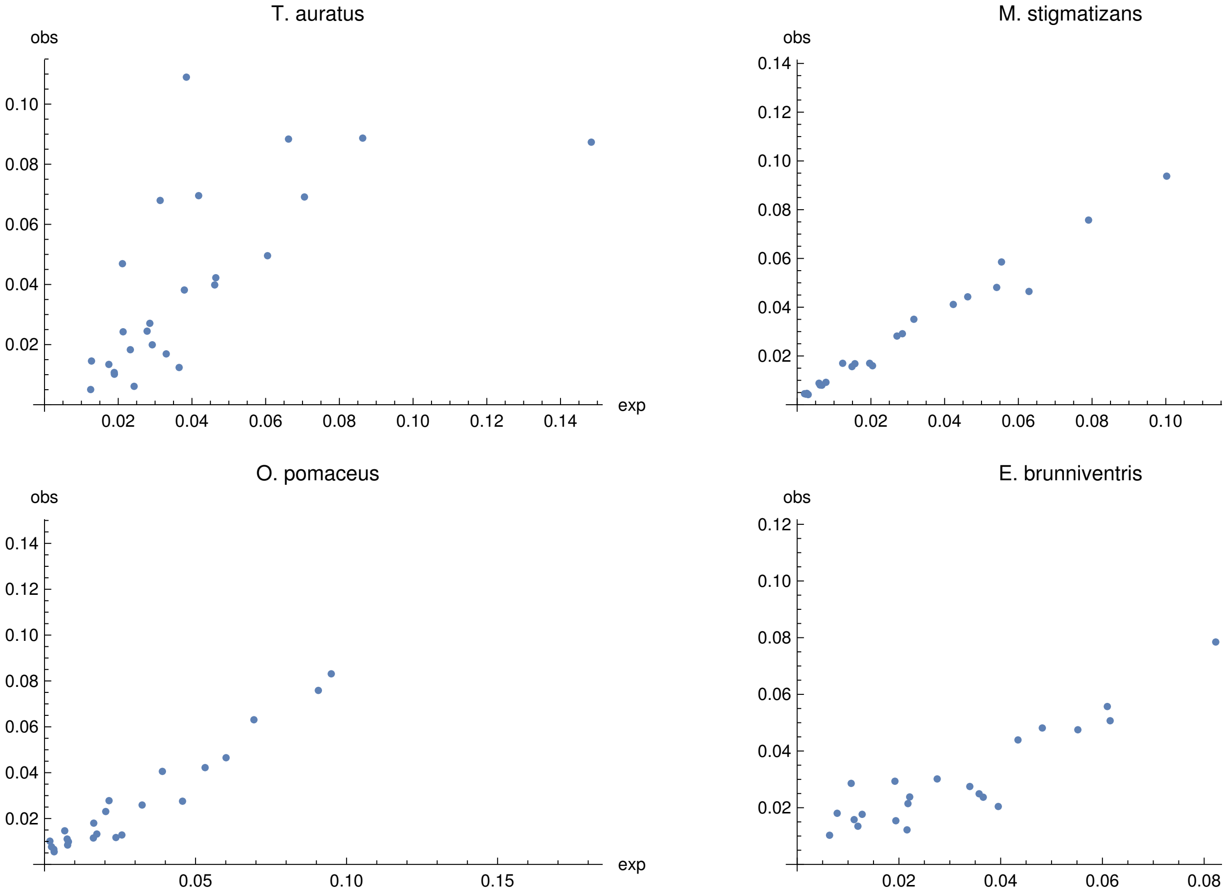


In[∘]:=
```
charts =
  Flatten[Table[{ToString[i], ToString[k]}, {i, 0, 4}, {k, 0, 4}] /. "4" → ">3", 1];
```

In[∘]:=
```
expbSFSnull = Table[likFullTab[Step5GFsolEInv // Total,
    ω, testnull⟦i, 2, 1, 2⟧, {λ1 → 1, T1 → 0}, 3], {i, 1, 7}];
expbSFSnullflat = Table[expbSFSnull⟦i⟧ // Normal // Thread // Flatten, {i, 1, 7}];
```

```
lblStyle = Style[#, 5, Black] &;

exptlnLBest = SetPrecision[
   Join[Table[allStep⟦i, 1⟧ / Total[(allcounts⟦i⟧ // Normal // Flatten)], {i, 1, 4}],
    Table[testnull⟦i, 1⟧ / Total[(allcounts⟦i⟧ // Normal // Flatten)], {i, 5, 7}]], 3];
```

In[•]:=
```
bigfig =
  GraphicsGrid[{Table[PairedBarChart[obsbSFSflat⟦i⟧ , expbSFSflat⟦i⟧, AxesLabel →
       {"freq", "bSFS"}, ChartLabels → lblStyle /@ (ToString[#] & /@ charts),
      ChartStyle → {{Orange, Black}, None, None}, LabelStyle → Black,
      PlotLabel → allSp⟦i⟧, Epilog → {Text[Style[StringJoin["lnCL=",
           ToString[exptlnLBest⟦i⟧]], 16], Scaled[{.08, .9}], {0, 1}]}], {i, 1, 4}],
    Table[PairedBarChart[obsbSFSflat⟦i⟧, expbSFSnullflat⟦i⟧, AxesLabel →
       {"freq", "bSFS"}, ChartLabels → lblStyle /@ (ToString[#] & /@ charts),
      ChartStyle → {{Orange, Gray}, None, None},
      LabelStyle → Black, PlotLabel → allSp⟦i⟧,
      Epilog → {Text[Style[StringJoin["lnCL=", ToString[exptlnLBest⟦i⟧]], 16],
         Scaled[{.08, .9}], {0, 1}]}], {i, 5, 7}]}, Spacings → 0, ImageSize → 1600]
```

Out[•]=

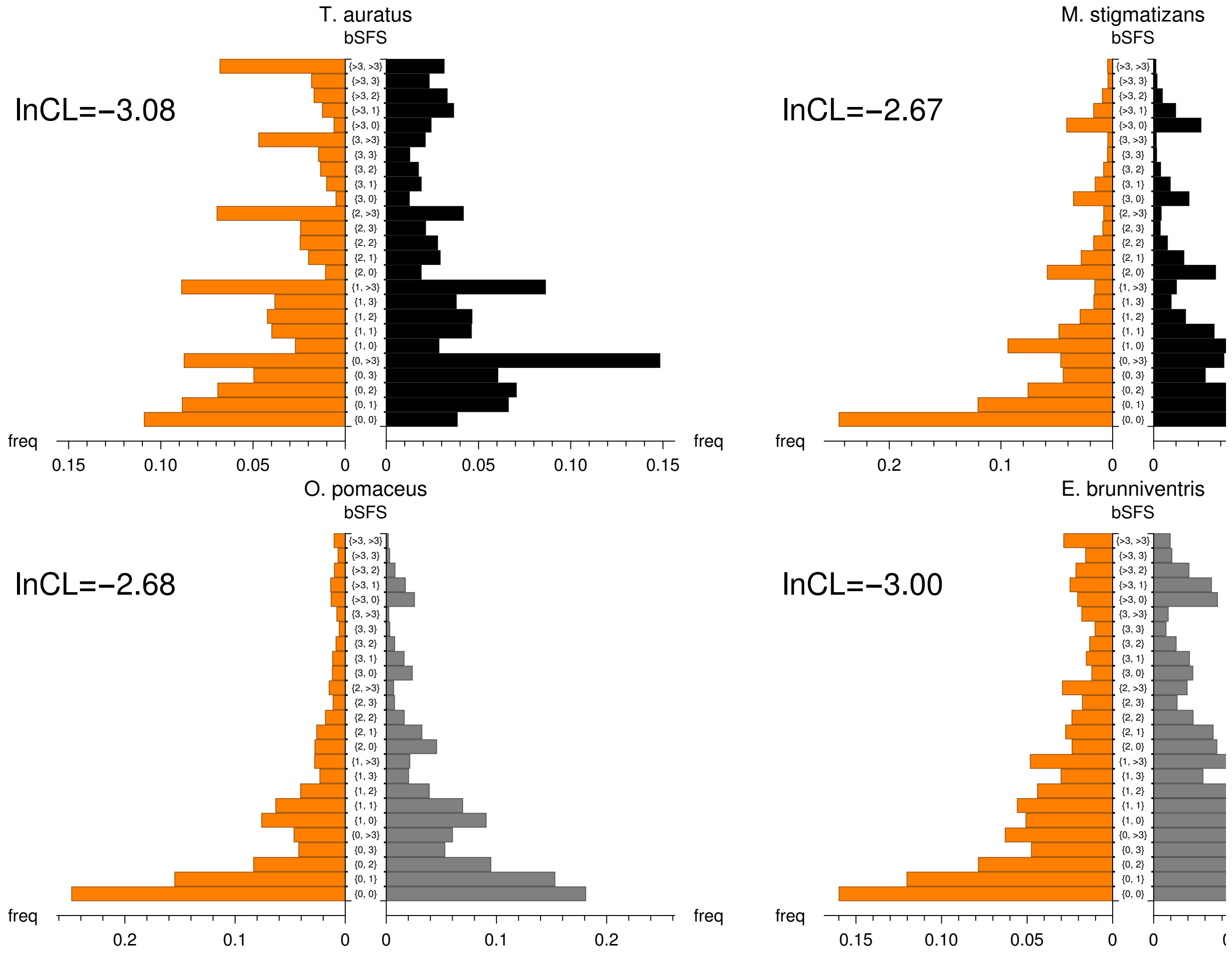

In[•]:=
```
Export[
 "/home/konrad/Dropbox/Manuscripts/Walton_Iberian_demography_2019/Figures/goodness
   .jpg", bigfig]
```

Out[•]=
```
/home/konrad/Dropbox/Manuscripts/Walton_Iberian_demography_2019/Figures/goodness.
 jpg
```

### Scaling parameter estimates

$N_{e1}$ is:

In[217]:=
```
ne1 = Table[allStep〚i, 2, 1, 2〛 / (allblockl〚i〛 * 4 * 3.46 × 10^-9), {i, 1, 7}];
ne2 = Table[ne1〚i〛 / allStep〚i, 2, 2, 2〛, {i, 1, 7}];
```

In[•]:=
```
neNull = Table[testnull〚i, 2, 1, 2〛 / (allblockl〚i〛 * 4 * 3.46 × 10^-9), {i, 1, 7}]

nepi = Table[piall〚i〛 / (4 * 3.46 × 10^-9), {i, 1, 7}];
```

In[219]:=
```
gen = {2, 1, 2, 2, 2, 2, 2};
```

In[220]:=
```
tEst = Table[allStep〚i, 2, 3, 2〛 * 2 ne1〚i〛 / 1000, {i, 1, 7}] / gen;
```

In[221]:=
```
SetPrecision[{allSp, ne1 / 10 000, ne2 / 10 000, tEst}, 5] // TableForm
```

Out[221]//TableForm=

| | | | | | |
|---|---|---|---|---|---|
| T. auratus | M. stigmatizans | M. dorsalis | O. nitidulus | O. pomaceus | E. bru |
| 23.854 | 1.7721 | 6.6588 | 4.7162 | 20.730 | 51.522 |
| 78.547 | 6.0670 | 14.395 | 19.432 | 20.183 | 3758.5 |
| 834.88 | 9.9652 | 12.467 | 33.148 | 1801.6 | 1567.6 |

### Relative maximum $N_e$

In[222]:=
```
rawπ = {0.0064, 0.00067, 0.0018, 0.0016, 0.0028, 0.0071, 0.0032};

peakPSMC = {83.5, 6.5, 13.9, 15.7, 27.5, 83.1, 21.3} * 10 000;

neraw = rawπ / (4 * 3.46 × 10^-9);
```

In[246]:=
```
{allSp, (Max[#] & /@ ({ne1, ne2} // Thread)) / nepi, peakPSMC / neraw} // Thread //
 TableForm
```

Out[246]//TableForm=

| | | |
|---|---|---|
| T. auratus | 2.13543 | 1.80569 |
| M. stigmatizans | 1.5521 | 1.34269 |
| M. dorsalis | 1.39703 | 1.06876 |
| O. nitidulus | 2.12689 | 1.35805 |
| O. pomaceus | 1.2467 | 1.35929 |
| E. brunniventris | 91.4543 | 1.61986 |
| C. fungosa | 4.09389 | 0.921225 |

### Definitions

In[202]:=

```
pi[l_] :=
  Module[{tsfs = Total[l]}, (tsfs⟦1⟧ 2 (1 / 5) (4 / 5) + tsfs⟦2⟧ 2 (2 / 5) (3 / 5))] // N;
```

In[203]:=

```
sfsnorm[l_] := Total[l] / Total[Total[l]] // N;
```
